# Impact of Molecular Crowding on Accessibility of Telomeric Overhangs Forming Multiple G-quadruplexes

**DOI:** 10.1101/2025.05.26.656241

**Authors:** Golam Mustafa, Sajad Shiekh, Janan Alfehaid, Sineth G. Kodikara, Hamza Balci

## Abstract

Telomeric overhangs, composed of repeating GGGTTA sequences, can fold into multiple G-quadruplex (GQ) structures that are essential for maintaining genomic stability and regulating telomere length. Molecular crowding--a defining feature of the cellular environment--affects folding kinetics, conformation, and stability of individual GQs. However, its influence on the overall architecture and accessibility of telomeric overhangs containing multiple GQs remains largely unexplored. In this study, we employed single-molecule Förster Resonance Energy Transfer (smFRET) and FRET-Point Accumulation for Imaging in Nanoscale Topography (FRET-PAINT) to address this question. We examined the accessibility of telomeric overhangs capable of forming one to six GQs to a short complementary peptide nucleic acid (PNA) imager probe under molecular crowding conditions. These conditions were simulated by polyethylene glycol (PEG) molecules of two different molecular weights: 200 Da (PEG-200) and 6000 Da (PEG-6000). Our results reveal a systematic reduction in the overhang accessibility with increasing PEG concentration—showing approximately a 3-fold reduction at 30% (v/v) PEG-200 and an 8-fold reduction at 30% PEG-6000. We also observed a progressive compaction of the overhang as PEG concentration increased, suggesting molecular crowding promotes architectural condensation, thereby reducing probe accessibility. These findings offer new insights into how the crowded cellular environment may compact telomeric overhangs and modulate their structural and functional properties.

## Introduction

The ends of human chromosomes, known as telomeres, consist of tandem repeats of the sequence GGGTTA (referred to as a ‘G-Tract’) and their complementary strands, extending approximately 10,000 base-pairs ^1–3^. Additionally, telomeres feature a 50-300 nucleotide (nt) long single-stranded G-rich overhang at the 3′-end. Four repeats of the GGGTTA sequence in the overhang can fold into a G-quadruplex (GQ) structure, composed of stacked G-quartets with four coplanar guanines ^4–6^. Multiple tandem GQs—ranging from 2 to 12-can assemble within the overhang, serving to protect chromosome ends by preventing their recognition as DNA double-strand breaks and safeguarding against end-to-end fusion and fraying ^7–9^. Additionally, GQ formation inhibits telomerase activity and contributes to the regulation of telomere length ^10^.

Environmental factors, such as monovalent cations and molecular crowding, play important roles in determining the stability, kinetics, structure, and polymorphism of individual GQ structures ^6,11–13^. However, despite recent efforts ^14–20^, important structural and functional questions regarding telomeric overhangs that contain multiple GQ structures remain largely unexplored ^21^. Notably, most *in vitro* studies on relevant systems have been performed in dilute, homogeneous buffer conditions. In contrast, the intracellular environment differs markedly from these buffer conditions, characterized by high concentrations of both soluble and insoluble biomolecules ^22–24^. Up to 40% of biomolecules in living cells cause molecular crowding, which is one of the distinctive characteristics of intracellular environment ^25^. Molecular crowding agents, along with associated changes in the hydration levels, significantly influence the topology of human telomeric GQ, inducing a conformational change from antiparallel to parallel ^26,27^. This transition has important implications for development of drugs targeting telomeric GQs ^7^. Furthermore, the stability of GQs and the rate of GQ formation increase in the presence of molecular crowding agents ^13,28,29^, which is significant in the context of telomerase-catalyzed telomere extension. Therefore, studying telomeric overhangs that can form multiple GQs under molecular crowding conditions can provide important insights about their physiological structure and function.

Using single molecule Förster Resonance Energy Transfer (smFRET) and FRET-Point Accumulation for Imaging in Nanoscale Topography (FRET-PAINT) ^30–32^, we investigated the accessibility of telomeric overhangs that contain 4-24 G-Tracts in the presence of a widely used molecular crowder polyethylene glycol (PEG), with molecular weights of 200 Da (PEG-200) and 6000 Da (PEG-6000). These telomeric overhangs can form 1-6 tandem GQs when fully folded; however, the number of folded GQs and their distribution depends on the folding pattern. By quantifying their accessibility to a short complementary strand, we investigated the impact of PEG-200 and PEG-6000 on the telomeric overhang architecture.

## Materials and Methods

### Nucleic Acid Constructs

The DNA strands were obtained from Eurofins Genomics (Louisville, KY, USA) or Integrated DNA Technologies (Coralville, IA, USA). The oligonucleotides were purified in-house using denaturing polyacrylamide gel electrophoresis (PAGE) ^17^. The HPLC purified short Cy5-PNA strand was purchased from PNA Bio (Thousand Oaks, CA, USA). The sequences of the oligos are as follows:

*nG-Tract*: TGGCGACGGCAGCGAGGC TTA (GGG TTA)_n_ G, where n represents the number of G-Tracts in the overhang (n=4, 6, 8…20, 22, 24).

*Stem-18-Cy3*: Cy3-GCCTCGCTGCCGTCGCCA-Biotin

*Cy5-PNA*: <u>TAA CCC T</u>T-Cy5

*16G-Tract-12nt*: TGGCGACGGCAGCGAGGC TT (GGGTTA)_16_ <u>GTACGATCGCAG</u>

*Stem-18-Cy5:* Cy5-GCCTCGCTGCCGTCGCCA-Biotin

*Cy3-DNA*: <u>CTGCGATCGTAC</u>-Cy3

*T30:* TGGCGACGGCAGCGAGGC T_30_-Cy3

The underlined segments of Cy3-DNA are complementary to the underlined segments of 16G-Tract-12nt, while the underlined segments of Cy5-PNA are complementary to the telomeric repeats. The Stem-18-Cy3 is complementary to the 18 nt segments at the 5’-ends of nG-Tract. Similarly, Stem-18-Cy5 strand is complementary to the 18 nt segments at the 5’-ends of 16G-Tract-12nt and T30 constructs. To mimic physiological telomeric structure, a partial-duplex DNA (pdDNA) construct with a double-stranded DNA (dsDNA) stem and single-stranded DNA (ssDNA) telomeric overhang was created. The pdDNA constructs were created by annealing a short stem strand (18 nt long) and a long strand (including a complementary sequence to the stem strand and an overhang). The annealing was performed in 1:5 ratio of short to long strand to ensure that all Stem-18 strands (biotinylated and Cy3 or Cy5-labeled) have a matching long strand. After the DNA is immobilized on the surface via the biotin-streptavidin linker, the unpaired long strands are washed out of the channel. The annealing reaction was performed in a thermal cycler (Hybaid Omn-E Thermal Cycler) at 150 mM KCl and 10 mM MgCl_2_ by heating the two strands at 95 °C for 3 minutes, then cooling to 30 °C in 1 °C steps with 3-minute incubation in each step. This slow cooling process ensures that the folding pattern achieves a thermodynamic steady state. After annealing, the MgCl_2_ concentration was reduced to 2 mM for all later stages of experiment.

### Sample preparation and smFRET and FRET-PAINT assays

The glass coverslips and laser-drilled quartz slides were thoroughly cleaned with acetone and potassium hydroxide (KOH), followed by piranha etching, aminosilane surface functionalization, and NHS-ester polyethylene glycol (PEG) surface passivation. A PEG mixture with a 100:4 ratio of m-PEG-5000:biotin-PEG-5000 (Laysan Bio Inc.) was used. PEG prevents non-specific binding of DNA and PNA molecules to the surface, whereas biotin-PEG-5000 provides attachment sites for biotinylated DNA molecules via streptavidin. To densify the PEG layer, a second round of PEGylation was performed with a small (333 Da) MS(PEG)_4_ (Thermo Fisher Scientific, Inc). The microfluidic sample chambers were created by sandwiching double-sided tape between a PEGylated slide and coverslip, followed by sealing the chamber with epoxy. The chamber was treated with 1% (v/v) Tween-20 for 15 minutes, followed by extensive washing and incubation of 0.01 mg/ml streptavidin for 2 minutes.

The pdDNA samples were diluted to 10-40 pM in 150 mM KCl and 2 mM MgCl_2_ before they were added to the chamber and incubated for 1-5 minutes, yielding a surface density of 250-350 molecules per imaging area (∼50 μm × 100 μm). Excess or unbound DNA was removed from the chamber by washing it with 150 mM KCl and 2 mM MgCl_2_. The imaging buffer contained 50 mM Tris-HCl (pH 7.5), 2 mM Trolox, 0.8 mg/mL glucose, 0.1 mg/mL glucose oxidase, 0.1 mg/mL bovine serum albumin (BSA), 2 mM MgCl_2_, 150 mM KCl, 40 nM Cy5-PNA and a specific concentration of PEG-200 or PEG-6000 over 0-30% (v/v) range. Prior to adding the Cy5-PNA strand stock solution to the imaging buffer, the solution was heated to 85 °C for 10 minutes to increase solubility. Prior to video recording, the imaging solution containing Cy5-PNA and PEG-200 (or PEG-6000) was incubated with the DNA molecules in the sample chamber for 15 minutes. In Fig. 2 and Fig. 3, we performed comparative analysis for 11 different DNA constructs at 0% and 15% PEG-200. In addition to being in the middle of the concentration range we studied, 15% PEG-200 also yielded high enough frequency of Cy5-PNA binding events, which is critical for attaining reliable statistics. Also, we observed a gradual degradation in imaging quality with an increasing PEG concentration. The 15% PEG-200 concentration was high enough to yield significantly different results compared to the absence of PEG-200, with a manageable decrease in binding event statistics and imaging quality.

**Fig. 1.**
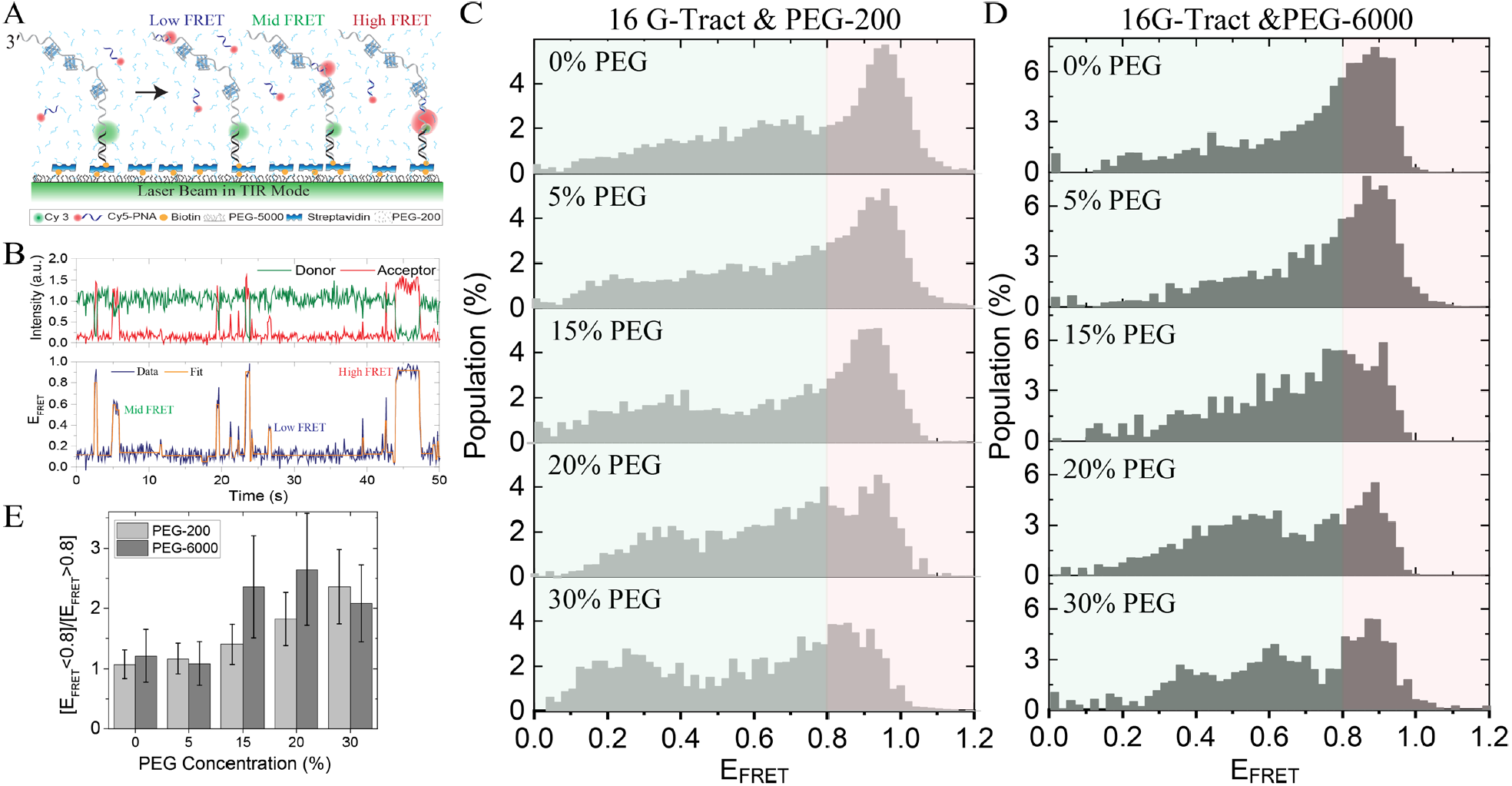
(A) Schematic of the FRET-PAINT assay. Donor (Cy3) labeled pdDNA constructs are immobilized on the surface via biotin-neutravidin attachment. The overhangs may form multiple GQs separated from each other by unfolded segments, which can be accessed by the complementary Cy5-PNA imager strand for a short period of time, resulting in pulses in acceptor intensity and E_FRET_. (B) An example FRET-PAINT time trace that shows binding to different segments of a single telomeric overhang results in different FRET levels, labeled with low, mid, and high FRET to highlight the differences--many more levels are observed over different molecules. The orange line is a fit to the FRET trace. Normalized FRET-PAINT histograms obtained from measurements on 16G-Tract construct in the presence of 0-30% (C) PEG-200 and (D) PEG-6000. The number of DNA molecules included in the analysis of each histogram is given in Supplementary Table S1. The average number for each histogram is 120 DNA molecules (a total 597 molecules) in Fig. 1C and 96 molecules (a total of 478 molecules) in Fig. 1D. (E) The ratio of the total population of [E_FRET_ < 0.8] to [E_FRET_ > 0.8], which is a measure of binding to the junction region. The error bars are standard error of the mean.

**Fig. 2.**
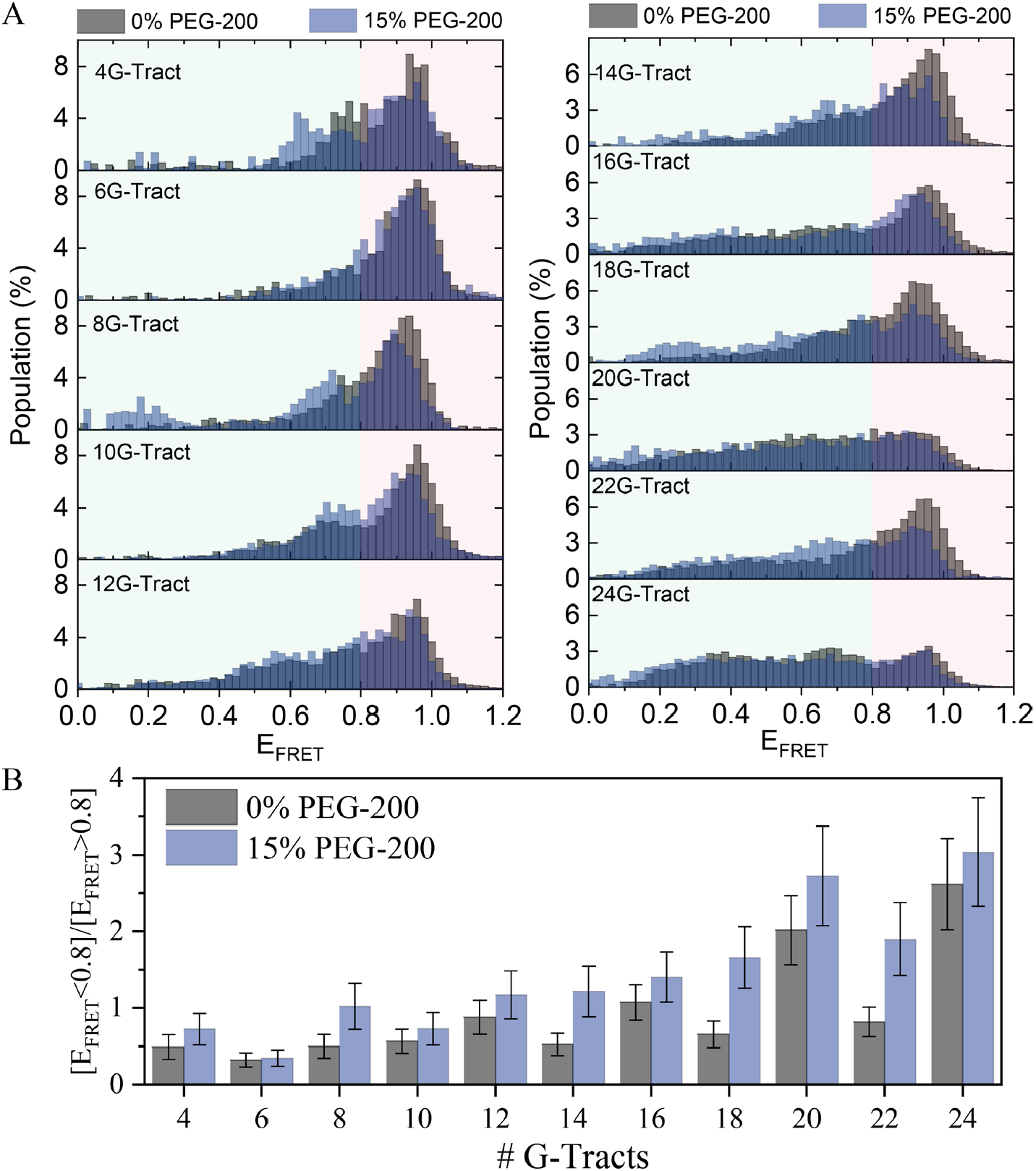
Normalized FRET-PAINT histograms in the absence and the presence of 15% PEG-200 for overhangs containing 4 to 24 G-Tracts. (B) The fraction of binding events to the junction region (E_FRET_>0.8) is generally smaller in the presence of 15% PEG-200, as quantified via the ratio of the total population of [E_FRET_ < 0.8] to that of [E_FRET_ > 0.8]. The error bars are standard error of the mean. The number of DNA molecules included in the analysis of each histogram is given in Supplementary Table S1, the average being 125 DNA molecules (a total 3119 molecules in Fig. 2A).

**Fig. 3.**
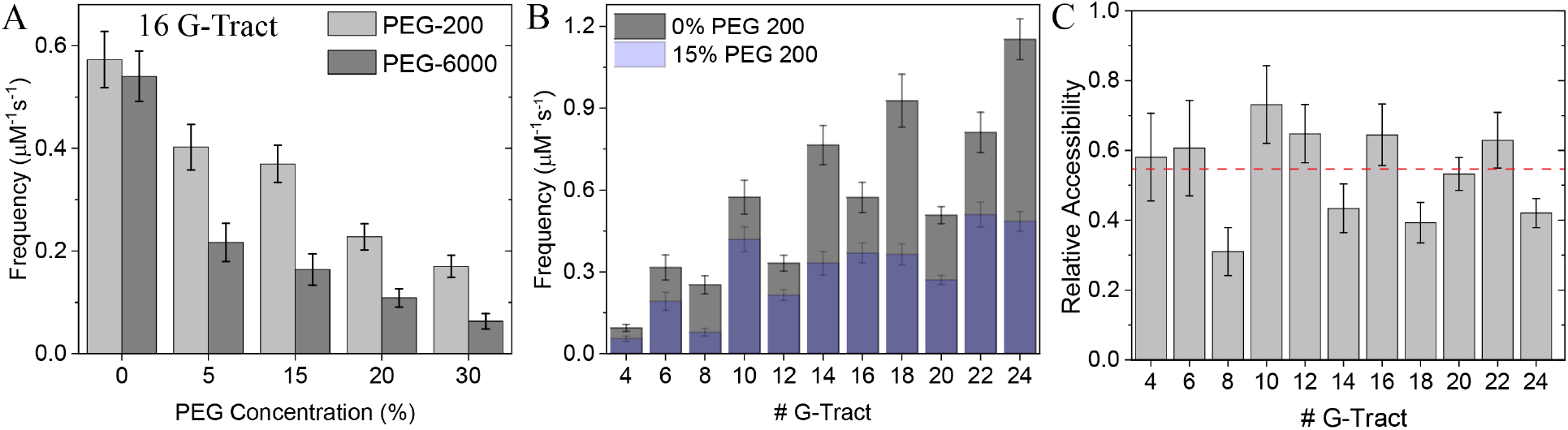
**A**nalysis of binding frequencies of Cy5-PNA to telomeric overhangs. The frequencies are normalized to Cy5-PNA concentration. (A) The binding frequency of Cy5-PNA to the 16 G-Tract construct decreases as the PEG-200 concentration increases in the 0-30% (v/v) range. (B) The binding frequency of Cy5-PNA to 4-24 G-Tract constructs in the absence and presence of 15% PEG-200. (C) The relative accessibility of 4-24G-Tract at 15% vs 0% PEG-200 is shown. When averaged over all constructs, the mean relative accessibility is 0.54±0.02 (red, dashed line). The error bars for the binding frequencies and the relative accessibility are calculated using a bootstrapping analysis (using 20000 different bootstrapping sets). The number of molecules for each histogram is given in Supplementary Table S2. On average, 580 DNA molecules (including those that did not show any Cy5-PNA binding) and 380 Cy5-PNA binding events were analyzed for each condition. A total of 14488 DNA molecules and 9504 Cy5-PNA binding events were analyzed in Fig. 3.

All measurements were carried out using a home-built prism-type total internal reflection fluorescence (TIRF) instrument equipped with an Olympus IX-71 microscope and an Andor IXON EMCCD camera (IXON DV-887 EMCCD, Andor Technology, CT, USA-part of Oxford Instruments). A total of 5-10 movies of 2000 frames per movie were recorded at a frame integration time of 100 ms/frame. A laser beam of 532 nm wavelength (Spectra Physics Excelsior) was used to excite the donor fluorophores. The fluorescence signal was collected using an Olympus water objective (60×, 1.20 NA).

### Data Analysis

Custom software written in C++ was used to generate single molecule time traces of donor (I_D_) and acceptor (I_A_) intensity from the recorded movies ^33^. The time traces of each molecule were screened using a custom MATLAB code to ensure they represent single molecules. Based on the remnant donor and acceptor intensities after donor photobleaching, the background was subtracted from each of these selected molecules. The FRET efficiency (E_FRET_) was calculated as E_FRET_ = I_A_/(I_A_ + I_D_). The screened single molecules that did not show any binding event contributed to a donor-only (DO) peak, which is due to leakage of donor emission into the acceptor channel. The DO peak was used as a reference for rescaling the FRET range and shifting the DO E_FRET_ to zero. Screened single molecule traces with at least one Cy5-PNA strand binding event were used to create smFRET or FRET-PAINT histograms.

The E_FRET_ levels in the FRET-PAINT time trace along with the AutoStepfinder fit (orange line) are shown in Fig. 1B ^34^. To distinguish a binding event from random signal fluctuations, only FRET pulses with dwell times greater than 300 ms and an E_FRET_ greater than 0.13 are included in the analysis as binding events. To find the binding frequency, the total number of binding events is divided by the total observation time. The number of binding events was counted using a custom MATLAB code that processed the fits generated by the AutoStepfinder program. The FRET-PAINT histograms were created from the raw time traces, rather than the idealized fits generated by AutoStepfinder.

## Results

In the FRET-PAINT measurements shown in Figs. 1-3, surface-immobilized DNA constructs were labeled with a donor fluorophore (Cy3) and served as the docking strands. A Cy5-labeled peptide nucleic acid (Cy5-PNA; sequence: <u>TAA CCC T</u>T-Cy5), complementary (underlined) to a 7-nt long telomeric sequence, was used as the imager strand. A schematic of the assay is shown in Fig. 1A. When Cy5-PNA binds to a G-Tract, a FRET signal is observed (Fig. 1B). This signal depends on the position of the binding site within the overhang and the folding pattern of the overhang, including distribution of the unfolded regions which serve as the primary binding sites for Cy5-PNA. A sample FRET-PAINT time trace demonstrating such binding events is shown in Fig. 1B, from which the frequency of binding events can be determined. The FRET-PAINT distributions (Fig. 1C-D and Fig. 2A) were created by combining the binding events in a histogram. The DNA constructs that do not show any Cy5-PNA binding, such as those that do not contain any unfolded or accessible sites, do not contribute to these histograms.

Fig. 1C and Fig. 1D show the FRET-PAINT histograms for the 16G-Tract construct (which can form up to four GQs) in the presence of PEG-200 and PEG-6000, respectively. As the PEG concentration increases, the relative contribution of Cy5-PNA binding events to the junction region (which results in E_FRET_ ≈ 0.9 since the Cy3 is at the junction) becomes less prominent while those of the intermediate regions and the 3′-end (which result in lower E_FRET_ as they are further from the Cy3) become more significant. To quantify this shift, we plotted the ratio of the cumulative binding events with E_FRET_<0.8 to those with E_FRET_>0.8 (Fig. 1C and Fig. 1D). The E_FRET_=0.8 level was selected as a reference to focus on the high FRET peak that represents binding to the vicinity of the junction. This ratio, [E_FRET_<0.8]/[E_FRET_>0.8], increases systematically with rising concentration of PEG-200 or PEG-6000 (Fig. 1E).

The FRET-PAINT distributions of telomeric overhangs ranging from 4G-Tract to 24G-Tract in the absence of PEG or the presence of 15% (v/v) PEG-200 are shown in Fig. 2A. At this PEG-200 concentration, the shift away from the junction region is of modest scale but a widespread feature, as illustrated by the [E_FRET_<0.8]/[E_FRET_>0.8] ratios in Fig. 2B. We also quantified the change in the broadness of the distributions for 0% and 15% PEG-200 cases, which showed only insignificant variations (Supplementary Fig. S1).

Fig. 3A shows the frequency of Cy5-PNA binding to the 16G-Tract construct in varying concentrations of PEG-200 or PEG-6000. The binding frequency systematically decreases as the concentration of either PEG is increased, with the decrease being significantly more prominent in the presence of PEG-6000 compared to PEG-200. At 30% concentration, PEG-200 and PEG-6000 induce approximately a 3-fold and an 8-fold reduction in the binding frequency compared to the absence of PEG, respectively. Fig. 3B shows comparative Cy5-PNA binding frequencies at 0% and 15% PEG-200 for 4-24G-Tract constructs. The effect of molecular crowding on telomeric accessibility can be quantified by calculating the relative accessibility, which is defined as the ratio of binding frequency in the presence of 15% PEG-200 to that in its absence (Fig. 3C). The mean relative accessibility obtained from all constructs is 0.54±0.02 (red, dashed line in Fig. 3C), suggesting on average a 2-fold drop at 15% PEG-200. Furthermore, in both 0% and 15% PEG-200 concentrations, the binding frequencies show a dip for overhangs with [4n] G-Tracts (e.g., 8, 12, 16) compared to those with [4n+2] G-Tracts (e.g., 10, 14, 18). The differences between these two groups in terms of folding frustration and steric hindrance have been discussed in an earlier publication ^17^. The two additional repeats in the [4n+2] G-Tract constructs ensure that at least two repeats are unfolded (n GQs require 4n G-Tracts) and accessible to Cy5-PNA, increasing the binding frequency.

Increasing the PEG concentration increases the viscosity of the environment, in addition to a modest increase in refractive index. In particular, the viscosity increases ∼3-fold in the presence of 30% PEG-200 compared to its absence ^35^. The change in viscosity impacts the diffusion of the imager strand Cy5-PNA and variations in the refractive index change the Förster radius of the FRET mechanism. In order to test the impact of such environmental factors, we performed FRET-PAINT experiments on a construct that contains a single binding site for Cy5-PNA. Experiments using both PEG-200 and PEG-6000 showed negligible variation in the FRET-PAINT distributions and the Cy5-PNA binding frequency (Supplementary Fig. S2). This suggests binding of Cy5-PNA to the overhang is not diffusion limited.

The reduction in Cy5-PNA binding frequency might have several reasons, including: (i) changes in the folding conformation of individual GQs; (ii) changes in the stability of individual GQs; and (iii) changes in the compactness of the overhang architecture. Changes in compactness could be driven by confinement of the unfolded regions to a smaller volume and by enhanced stacking interactions between neighboring GQs. These mechanisms are not mutually exclusive and could all take place to result in the observed reduction in accessibility. To illustrate, it is possible that in the presence of molecular crowders, GQs might attain uniform parallel conformation and higher stability, which might increase the probability of stacking with neighboring GQs, while the unfolded segments are sequestered into a smaller volume.

We propose that compaction of the overhang architecture, which reduces the accessibility of the unfolded segments, plays a significant role in decreasing the binding frequency. In an earlier study, we demonstrated that Cy5-PNA does not unfold the telomeric GQ at a significant level even in the absence of crowding agents. The thermal melting point of PNA-DNA duplex is T_m_ ≈ 12.6±0.3 °C ^17^, compared to T_m_ = 68 °C for unfolding the GQ in the absence of crowding agents ^36^. The Cy5-PNA binding frequency is about an order of magnitude greater for an overhang with a single G-Tract (one binding site) compared to that with four G-Tracts, which can fold into GQ ^17^. Therefore, compaction of the unfolded G-Tracts, which have much higher accessibility compared to those within GQ, should reduce the binding frequency of Cy5-PNA.

To directly test whether the overhang becomes more compact with increasing PEG concentration, we designed a telomeric DNA construct in which a single binding site for an imager strand is placed at the 3′-end of a 16G-Tract construct (Fig. 4A). The sequence of this binding site and that of the complementary imager strand are non-telomeric to ensure that the imager strand (labeled with Cy3) binds only to this site at the 3′-end. With the Cy5 fluorophore placed at the junction region (near 5′-end of the overhang), the binding events effectively measure the end-to-end separation of the overhang, hence its compactness. Fig. 4B shows these data in which the distributions systematically shift to higher FRET levels as the concentration of PEG-200 is increased in the 0-30% range, providing strong support for compaction of the overhang.

**Fig. 4.**
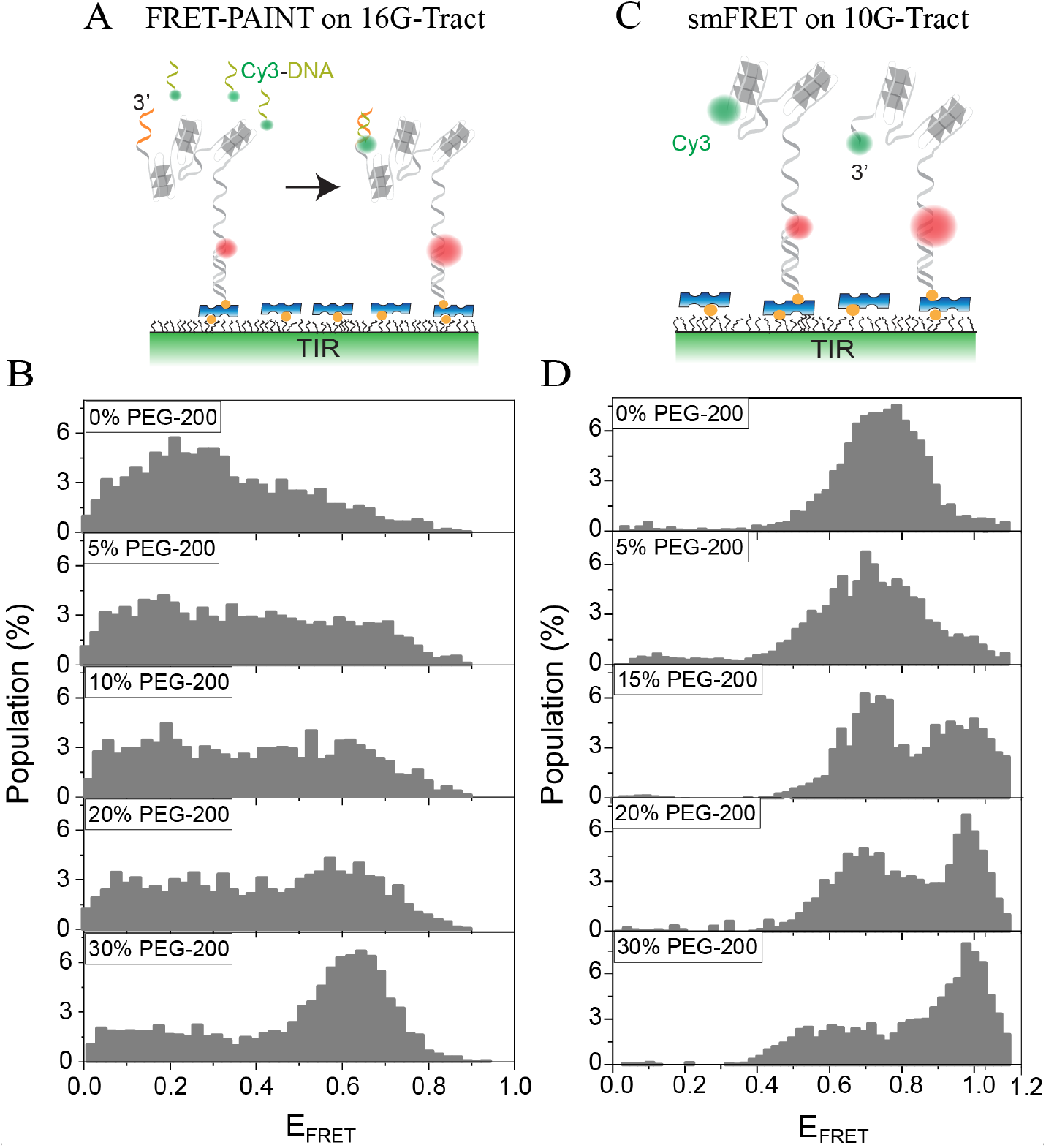
FRET-PAINT measurements on the 16G-Tract and smFRET measurements on the 10G-Tract construct. Both assays probe compaction of the telomeric overhang with increasing PEG-200 concentration. (A) Schematic of the FRET-PAINT assay where a single binding site (orange segment) for the imager probe (Cy3-DNA) is placed at the 3′-end of the overhang. (B) FRET-PAINT histograms illustrate a shift towards higher E_FRET_ as the PEG-200 concentration is increased, suggesting a compaction of the overhang. (C) Schematic of the smFRET assay where Cy3 is at the free 3′-end and Cy5 at the junction. (D) In agreement with the FRET-PAINT measurements, FRET histograms also illustrate a shift towards higher E_FRET_ as the PEG-200 concentration is increased.

We also performed smFRET experiments on a 10G-Tract construct which was labeled with a Cy3 molecule at the 3′-end of the overhang and a Cy5 at the junction region (Fig. 4C-D). In agreement with the FRET-PAINT studies on the 16G-Tract construct, the smFRET studies on the 10G-Tract construct also showed a systematic shift to upper FRET values with increasing PEG-200 concentration, suggesting the distance between the ends of the overhang decreases with increasing PEG-200 concentration.

To investigate whether the unfolded segments of the overhang are compacted with increasing PEG concentration, we performed smFRET experiments on a DNA construct (pd-T30) that has a 30 thymine overhang, which is not expected to form any secondary structures (Supplementary Figure S3). These studies also showed an upward FRET shift with increasing PEG-200 concentration. Finally, we performed FRET-PAINT studies on an overhang that has a single G-Tract that is separated from the junction region by a 10T unstructured region (Supplementary Figure S4). In this design, the unstructured region is significantly shorter and mimics the length when a G-Tract is not unfolded (9 nt). This construct also demonstrates a modest, yet detectable upward shift in the FRET-PAINT distribution with increasing PEG-200 concentration, which suggests the compaction of even a single unfolded G-Tract yields detectable compaction in the FRET distributions.

The crowding-induced compaction of the telomeric overhang raises questions about how their interactions with telomere-specific proteins or small molecules would be impacted. While a more stable GQ structure might facilitate the binding of small molecules ^37^, any compaction in the overhang might result in reduced accessibility for the molecules. Similarly, while excluded volume effects or dehydration might increase local concentration of proteins and promote interactions with telomeres, a more compact architecture might reduce access to the overhang. Further research is needed to address these interesting questions.

## Conclusions

In this study, we investigated the influence of crowding agents on the accessibility and architecture of telomeric overhangs ranging in length from 4 to 24 G-Tracts under dilute and crowded conditions. We observed that as the concentration of molecular crowding agent increases, the accessibility to telomeric overhang decreases by approximately 3-fold in 30% PEG-200 and approximately 8-fold in 30% PEG-6000. We demonstrate that compaction of the overhang, which better shields the unfolded segments that act as binding sites for the imager probe, plays a significant role in reducing accessibility. We also found that the fraction of binding events to the junction region, which is otherwise the most accessible region of the overhang, is reduced in the presence of crowding agents. Given the importance of telomeric overhang accessibility and compaction in maintaining genomic integrity, our study highlights the critical role the crowded cellular environment may play in stabilizing and compacting these vulnerable genomic regions.

### Supporting Information Available

Quantification of the broadness of FRET-PAINT distributions, frequency of Cy5-PNA binding events for different control experiments, and number of molecules and binding events included in each figure.

## Acknowledgements

This study was supported by the National Institutes of Health under Grant No. 1R35GM156183 and 1R15GM146180.

## Notes

### Competing Interest Statement

The authors have declared no competing interest.

## References

(1) Wright, W. E.; Tesmer, V. M.; Huffman, K. E.; Levene, S. D.; Shay, J. W. Normal Human Chromosomes Have Long G-Rich Telomeric Overhangs at One End. 1997, 11 (21), 2801–2809. 10.1101/gad.11.21.2801.

(2) Lim, C. J.; Cech, T. R. Shaping Human Telomeres: From Shelterin and CST Complexes to Telomeric Chromatin Organization; Nature Reviews Molecular Cell Biology, 2021; Vol. 22, pp 283–298. 10.1038/S41580-021-00328-Y.

(3) Blackburn, E. H.; E.H., B. Structure and Function of Telomeres. Nature 1991, 350 (6319), 569–573. 10.1038/350569a0.

(4) Biffi, G.; Tannahill, D.; McCafferty, J.; Balasubramanian, S. Quantitative Visualization of DNA G-Quadruplex Structures in Human Cells. Nat Chem 2013, 5 (3), 182–186. 10.1038/nchem.1548.

(5) Phan, A. T.; Mergny, J. L. Human Telomeric DNA: G-Quadruplex, i-Motif and Watson-Crick Double Helix. Nucleic Acids Res 2002, 30 (21), 4618–4625. 10.1093/nar/gkf597.

(6) Tran, P. L.; Mergny, J. L.; Alberti, P. Stability of Telomeric G-Quadruplexes. Nucleic Acids Res 2011, 39 (8), 3282–3294. https://doi.org/gkq1292[pii] 10.1093/nar/gkq1292.

(7) Hänsel-Hertsch, R.; Di Antonio, M.; Balasubramanian, S. DNA G-Quadruplexes in the Human Genome: Detection, Functions and Therapeutic Potential. Nat Rev Mol Cell Biol 2017, 18 (5), 279–284. 10.1038/NRM.2017.3.

(8) Yu, H. Q.; Miyoshi, D.; Sugimoto, N. Characterization of Structure and Stability of Long Telomeric DNA G-Quadruplexes. 2006, 128 (48), 15461–15468. 10.1021/ja064536h.

(9) Mendez-bermudez, A.; Hills, M.; Pickett, H. A.; Phan, A. T.; Mergny, J. L.; Riou, J. F.; Royle, N. J. Human Telomeres That Contain (CTAGGG)n Repeats Show Replication Dependent Instability in Somatic Cells and the Male Germline. Nucleic Acids Res 2009, 37 (18), 6225–6238. 10.1093/nar/gkp629.

(10) Zaug, A. J.; Podell, E. R.; Cech, T. R. Human POT1 Disrupts Telomeric G-Quadruplexes Allowing Telomerase Extension in Vitro. Proc Natl Acad Sci U S A 2005, 102 (31), 10864–10869. 10.1073/pnas.0504744102.

(11) Phan, A. T.; Luu, K. N.; Patel, D. J. Different Loop Arrangements of Intramolecular Human Telomeric (3+1) G-Quadruplexes in K+ Solution. Nucleic Acids Res 2006, 34 (19), 5715–5719. 10.1093/nar/gkl726.

(12) Chaires, J. B. Human Telomeric G-Quadruplex: Thermodynamic and Kinetic Studies of Telomeric Quadruplex Stability. FEBS J 2010, 277 (5), 1098. 10.1111/J.1742-4658.2009.07462.X.

(13) Aznauryan, M.; Birkedal, V. Dynamics of G-Quadruplex Formation under Molecular Crowding. Journal of Physical Chemistry Letters 2023, 14 (46), 10354–10360. 10.1021/ACS.JPCLETT.3C02453.

(14) Mustafa, G.; Shiekh, S.; Gc, K.; Abeysirigunawardena, S.; Balci, H. Interrogating Accessibility of Telomeric Sequences with FRET-PAINT: Evidence for Length-Dependent Telomere Compaction. Nucleic Acids Res 2021, 49 (6), 3371–3380. 10.1093/nar/gkab067.

(15) Abraham Punnoose, J.; Ma, Y.; Hoque, M. E.; Cui, Y.; Sasaki, S.; Guo, A. H.; Nagasawa, K.; Mao, H. Random Formation of G-Quadruplexes in the Full-Length Human Telomere Overhangs Leads to a Kinetic Folding Pattern with Targetable Vacant G-Tracts. Biochemistry 2018, 57 (51), 6946–6955. 10.1021/acs.biochem.8b00957.

(16) Yu, H.; Gu, X.; Nakano, S. I.; Miyoshi, D.; Sugimoto, N. Beads-on-a-String Structure of Long Telomeric Dnas under Molecular Crowding Conditions. J Am Chem Soc 2012, 134 (49), 20060–20069.

(17) Shiekh, S.; Mustafa, G.; Kodikara, S. G.; Hoque, M. E.; Yokie, E.; Portman, J. J.; Balci, H. Emerging Accessibility Patterns in Long Telomeric Overhangs. Proc Natl Acad Sci U S A 2022, 119 (30). 10.1073/pnas.2202317119.

(18) Shiekh, S.; Jack, A.; Saurabh, A.; Mustafa, G.; Kodikara, S. G.; Gyawali, P.; Hoque, M. E.; Pressé, S.; Yildiz, A.; Balci, H. Shelterin Reduces the Accessibility of Telomeric Overhangs. Nucleic Acids Res 2022, 50 (22), 12885–12895. 10.1093/NAR/GKAC1176.

(19) Monsen, R. C.; Trent, J. O.; Chaires, J. B. G-Quadruplex DNA: A Longer Story. Acc Chem Res 2022, 2022, 54. 10.1021/acs.accounts.2c00519.

(20) Monsen, R. C.; Chakravarthy, S.; Dean, W. L.; Chaires, J. B.; Trent, J. O. The Solution Structures of Higher-Order Human Telomere G-Quadruplex Multimers. Nucleic Acids Res 2021, 49 (3), 1749–1768. 10.1093/NAR/GKAA1285.

(21) Shiekh, S.; Kodikara, S. G.; Balci, H. Structure, Topology, and Stability of Multiple G-Quadruplexes in Long Telomeric Overhangs. J Mol Biol 2023, 168205. 10.1016/J.JMB.2023.168205.

(22) Zimmerman, S. B.; Trach, S. O. Estimation of Macromolecule Concentrations and Excluded Volume Effects for the Cytoplasm of Escherichia Coli. J Mol Biol 1991, 222 (3), 599–620. 10.1016/0022-2836(91)90499-V.

(23) Minton, A. P. The Effect of Volume Occupancy upon the Thermodynamic Activity of Proteins: Some Biochemical Consequences. Mol Cell Biochem 1983, 55 (2), 119–140. 10.1007/BF00673707.

(24) Ellis, R. J.; Minton, A. P. Join the Crowd. Nature. Nature Publishing Group September 2003, pp 27–28. 10.1038/425027a.

(25) Ellis, R. J. Macromolecular Crowding: Obvious but Underappreciated. Trends in Biochemical Sciences. Elsevier Current Trends October 2001, pp 597–604. 10.1016/S0968-0004(01)01938-7.

(26) Xue, Y.; Kan, Z. Y.; Wang, Q.; Yao, Y.; Liu, J.; Hao, Y. H.; Tan, Z. Human Telomeric DNA Forms Parallel-Stranded Intramolecular G-Quadruplex in K+ Solution under Molecular Crowding Condition. J Am Chem Soc 2007, 129 (36), 11185–11191. 10.1021/ja0730462.

(27) Miyoshi, D.; Nakao, A.; Sugimoto, N. Molecular Crowding Regulates the Structural Switch of the DNA G-Quadruplex. Biochemistry 2002, 41 (50), 15017–15024. 10.1021/bi020412f.

(28) Miller, M. C.; Buscaglia, R.; Chaires, J. B.; Lane, A. N.; Trent, J. O. Hydration Is a Major Determinant of the G-Quadruplex Stability and Conformation of the Human Telomere 3’ Sequence of d(AG3(TTAG3)3). J Am Chem Soc 2010, 132 (48), 17105–17107. 10.1021/ja105259m.

(29) Miyoshi, D.; Karimata, H.; Sugimoto, N. Hydration Regulates Thermodynamics of G-Quadruplex Formation under Molecular Crowding Conditions. J Am Chem Soc 2006, 128 (24), 7957–7963. 10.1021/ja061267m.

(30) Kodikara, S. G.; Merkel, K. J.; Haas, S. J.; Shiekh, S.; Balci, H. Detecting Secondary Structure Formation with FRET-PAINT. 2023. 10.1039/d3an01118f.

(31) Filius, M.; Kim, S. H.; Severins, I.; Joo, C. High-Resolution Single-Molecule FRET via DNA EXchange (FRET X). Nano Lett 2021, 21 (7), 3295–3301. 10.1021/acs.nanolett.1c00725.

(32) Auer, A.; Strauss, M. T.; Schlichthaerle, T.; Jungmann, R. Fast, Background-Free DNA-PAINT Imaging Using FRET-Based Probes. Nano Lett 2017, 17 (10), 6428–6434. 10.1021/acs.nanolett.7b03425.

(33) Lee, K. S.; Ha, T. SmCamera: All-in-One Software Package for Single-Molecule Data Acquisition and Data Analysis. Journal of the Korean Physical Society 2024, 86 (1), 1–13. 10.1007/S40042-024-01243-Z/TABLES/3.

(34) Loeff, L.; Kerssemakers, J. W. J.; Joo, C.; Dekker, C. AutoStepfinder: A Fast and Automated Step Detection Method for Single-Molecule Analysis. Patterns 2021, 2 (5). 10.1016/j.patter.2021.100256.

(35) Kirinčič, S.; Klofutar, C. Viscosity of Aqueous Solutions of Poly(Ethylene Glycol)s at 298.15 K. Fluid Phase Equilib 1999, 155 (2), 311–325. 10.1016/S0378-3812(99)00005-9.

(36) Qureshi, M. H.; Ray, S.; Sewell, A. L.; Basu, S.; Balci, H. Replication Protein A Unfolds G-Quadruplex Structures with Varying Degrees of Efficiency. Journal of Physical Chemistry B 2012, 116 (19), 5588–5594.

(37) Maleki, P.; Ma, Y.; Iida, K.; Nagasawa, K.; Balci, H. A Single Molecule Study of a Fluorescently Labeled Telomestatin Derivative and G-Quadruplex Interactions. Nucleic Acids Res 2017, 45 (1), 288–295. 10.1093/nar/gkw1090.

