## Supplementary Information for "Impact of Molecular Crowding on Accessibility of Telomeric Overhangs Forming Multiple G-quadruplexes"

### Quantifying the width of the FRET-PAINT Distributions

The FRET-PAINT histograms represent the distribution of Cy5-PNA binding sites to different segments of the overhang. Broader histograms suggest the accessible sites are distributed throughout the overhang while sharp histograms suggest the accessible sites are concentrated in specific regions. For example, if the junction region between the single and double stranded DNA is the most accessible region, the distributions will show a dominant high FRET peak since the donor fluorophore is located at that junction.

The broadness of the FRET-PAINT histograms is quantified by the S-parameter: $S = -\sum_i p_i ln\, p_i$ where the summation is carried over the entire FRET range and $p_i$ is the unfolded probability, normalized so that $\sum_{i=1}^{n} p_i = 1$ [1]. The S-parameter is analogous to the Shannon entropy of a specific FRET-PAINT distribution, with larger S-parameters indicating greater uncertainty in determining the position of the binding site. The FRET-PAINT histograms were normalized to a total of 100% in such a way that each molecule contributed equally to the histogram. Dividing this normalized FRET distribution by 100 gives the probability distribution for different FRET levels where the total probability is normalized to 1.0. More details about the S-parameter and its comparison with other statistical analysis methods are available in an earlier publication [1].

Figure S1 shows the S-parameter of the FRET-PAINT distributions shown in Figure 2. A small broadening is observed in most of the constructs. However, the observed effect is unanimously very small.

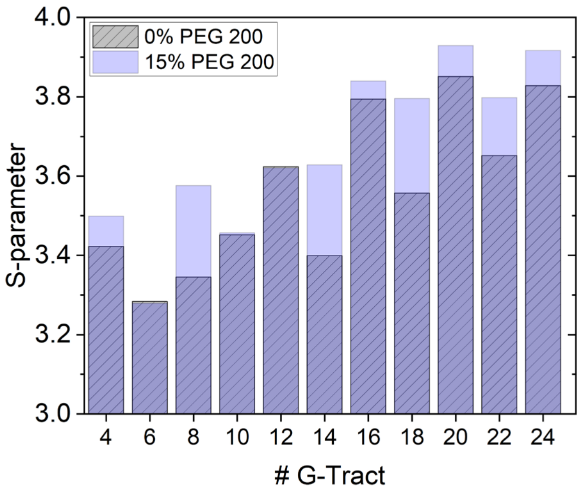


Figure S1. S-parameter calculations for the FRET-PAINT histograms shown in Figure 2A.

#### Testing the Impact of Viscosity on Imager Strand Binding Frequency

Fig. S2 shows FRET-PAINT experiments on a partial duplex DNA construct that has a single G-Tract at the overhang, which is not expected to form any secondary structure and serves as the single binding site for the imager strands Cy5-PNA (Fig. S2A-B). This simplified design allows directly measuring whether increased viscosity of the environment, when higher concentrations of PEG-200 are introduced, impacts the FRET-PAINT distributions and binding frequency of Cy5-PNA. As illustrated in Fig. S2C and Fig. S2D, the FRET-PAINT distributions and the binding frequencies, respectively, remain largely unchanged in the 0-30% PEG-200.

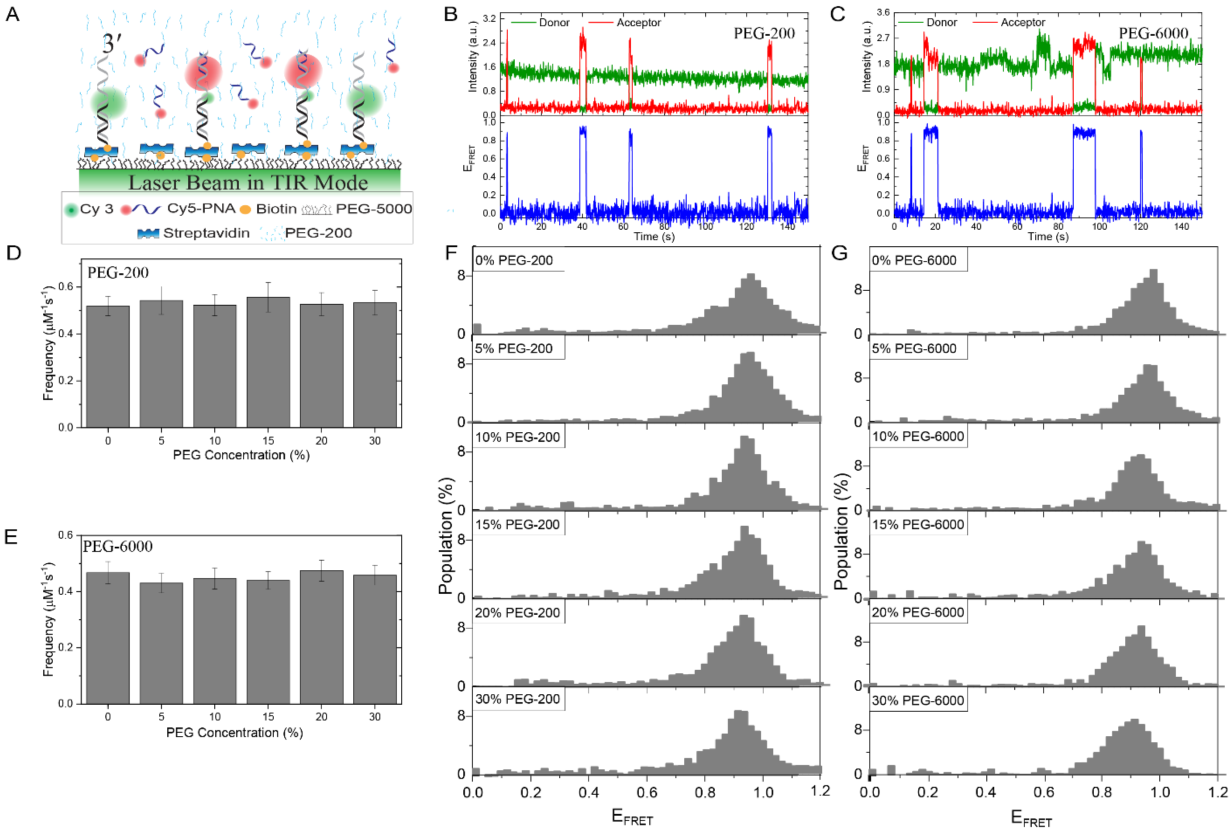


Figure S2. FRET-PAINT studies on the 1G-Tract construct to investigate the impact of viscosity on FRET-PAINT distributions and Cy5-PNA binding frequencies in the presence of PEG-200 or PEG-6000. (A) Schematic of the assay. (B) Sample FRET-PAINT time trace in PEG-200. (C) Sample FRET-PAINT time trace in PEG-6000. (D) Cy5-PNA binding frequencies in 0-30% PEG-200. (E) Cy5-PNA binding frequencies in 0-30% PEG-6000. (F) FRET-PAINT distributions in 0-30% PEG-200. (G) FRET-PAINT distributions in 0-30% PEG-6000. The number of molecules in each graph are given in Table S3.

### smFRET Measurements on a DNA Construct with Poly-Thymine Overhang

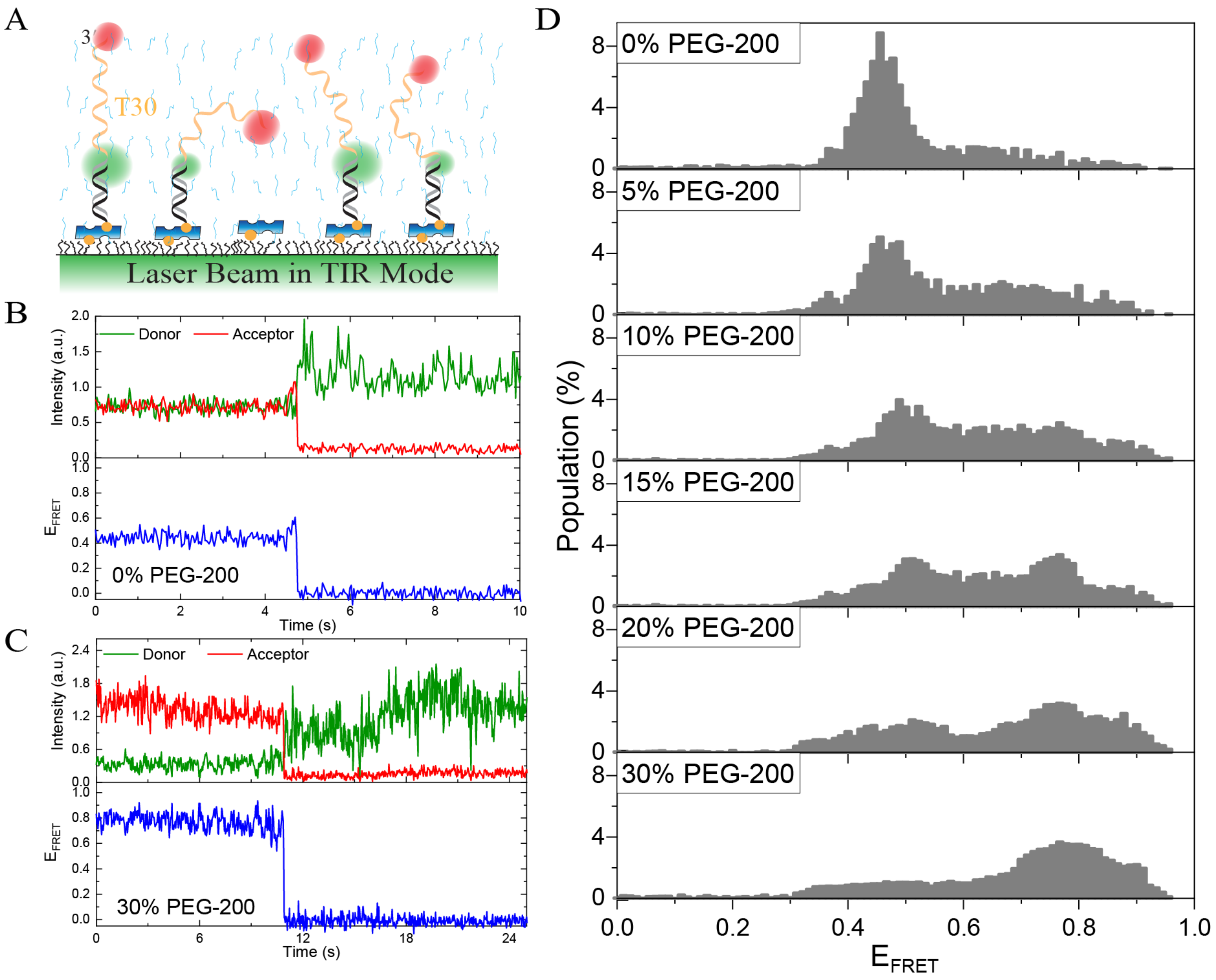


Figure S3. SmFRET measurements investigating compaction of an unstructured overhang, containing 30 thymine bases (pd-T30 construct). (A) Schematic of the smFRET assay. (B) A sample smFRET trace in the absence of PEG. (C) A sample smFRET trace in 30% PEG-200. (D) SmFRET histograms show gradual upward shift of the distributions to higher FRET values as PEG-200 concentration is increased.

**FRET-PAINT Measurements on a DNA Construct with 10T Spacer+1G-Tract Overhang**

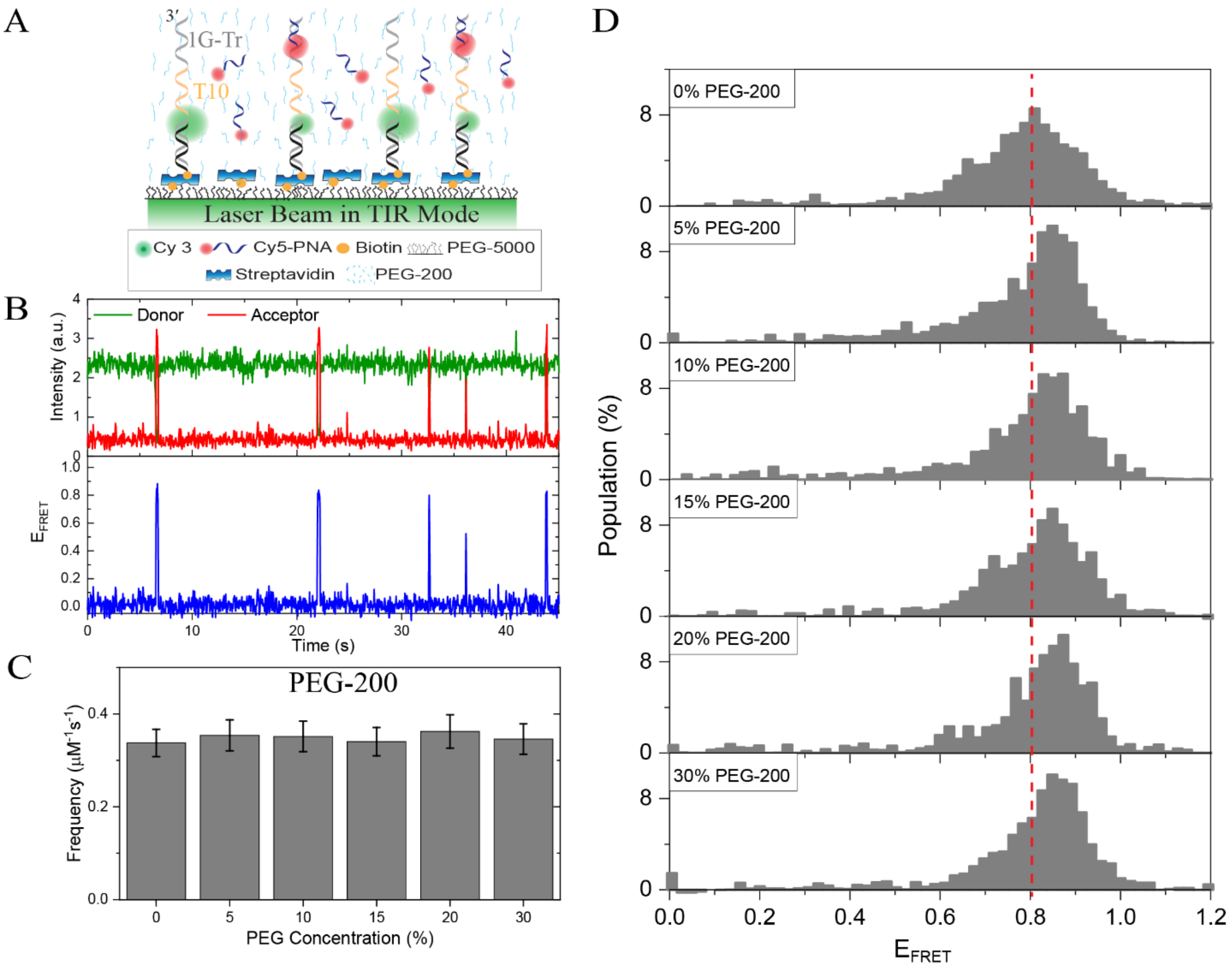


Figure S4. FRET-PAINT measurements investigating compaction of an overhang that contains 10 thymine bases between the duplex and the binding site for the imager strand (1G-Tr). (A) Schematic of the FRET-PAINT assay. (B) A sample FRET-PAINT trace. (C) Cy5-PNA binding frequencies in 0-30% PEG-200. (D) FRET-PAINT histograms showing gradual upward shift of the distributions to higher FRET values as PEG-200 concentration is increased. This increase suggests the 10T spacer between the binding site of the imager probe Cy5-PNA and the donor fluorophore is compacted as PEG-200 concentration is increased. The number of molecules in each graph are given in Table S3.

Table S1. Total number of DNA molecules ($N_{DNA}$) with at least one PNA binding event. PNA binding events to these molecules were used to construct the histograms in Fig. 1C-D and Figure 2A.

| Construct | [PEG] %(v/v) | $N_{DNA}$ |
|---|---|---|
| 4G-Tract | 0 | 67 |
| | 15 | 40 |
| 6G-Tract | 0 | 55 |
| | 15 | 44 |
| 8G-Tract | 0 | 81 |
| | 15 | 35 |
| 10G-Tract | 0 | 114 |
| | 15 | 87 |
| 12-Tract | 0 | 150 |
| | 15 | 134 |
| 14G-Tract | 0 | 130 |
| | 15 | 74 |
| 16G-Tract (PEG-200) | 0 | 163 |
| | 5 | 129 |
| | 15 | 131 |
| | 20 | 100 |
| | 30 | 74 |
| 16G-Tract (PEG-6000) | 0 | 147 |
| | 5 | 109 |
| | 15 | 106 |
| | 20 | 68 |
| | 30 | 48 |
| 18G-Tract | 0 | 104 |
| | 15 | 117 |
| 20G-Tract | 0 | 200 |
| | 15 | 156 |
| 22G-Tract | 0 | 155 |
| | 15 | 166 |
| 24G-Tract | 0 | 381 |
| | 15 | 232 |

Table S2. Statistics used to calculate binding frequency in Figure 3. $N_{Trace}$ refers to number of FRET time traces analyzed, $N_B$ to number of PNA binding events, $T_{tot}$ to total observation time, $f$ to binding frequency ($N_B/T_{tot}$), and $f_0/f_{15}$ to ratio of frequencies at 0 and 15% PEG-200.

| **Construct** | **[PEG] (v/v) %** | **$N_{trace}$** | **$N_B$** | **$T_{tot}$ (s)** | **f (×$10^{-3}$ $s^{-1}$)** | **$f_0/f_{15}$** |
|---|---|---|---|---|---|---|
| 4G-Tract | 0 | 812 | 134 | 56569 | 2.37 | 0.58 |
| | 15 | 715 | 77 | 55982 | 1.38 | |
| 6G-Tract | 0 | 411 | 158 | 19974 | 7.91 | 0.61 |
| | 15 | 478 | 135 | 28133 | 4.80 | |
| 8G-Tract | 0 | 496 | 229 | 36268 | 6.31 | 0.31 |
| | 15 | 763 | 73 | 37263 | 1.96 | |
| 10G-Tract | 0 | 382 | 388 | 27046 | 14.35 | 0.73 |
| | 15 | 374 | 251 | 23930 | 10.49 | |
| 12-Tract | 0 | 681 | 330 | 39817 | 8.29 | 0.65 |
| | 15 | 914 | 267 | 49713 | 5.37 | |
| 14G-Tract | 0 | 409 | 629 | 32911 | 19.11 | 0.43 |
| | 15 | 393 | 182 | 21939 | 8.30 | |
| 16G-Tract | 0 | 613 | 582 | 40632 | 14.32 | 0.65 |
| | 5 | 602 | 390 | 38758 | 10.06 | |
| | 15 | 581 | 309 | 33452 | 9.24 | |
| | 20 | 598 | 184 | 32297 | 5.70 | |
| | 30 | 657 | 125 | 29380 | 4.25 | |
| 18G-Tract | 0 | 231 | 485 | 20915 | 23.19 | 0.39 |
| | 15 | 579 | 308 | 33832 | 9.10 | |
| 20G-Tract | 0 | 509 | 512 | 40345 | 12.69 | 0.53 |
| | 15 | 472 | 251 | 37131 | 6.76 | |
| 22G-Tract | 0 | 473 | 752 | 37078 | 20.03 | 0.64 |
| | 15 | 656 | 530 | 41554 | 12.75 | |
| 24G-Tract | 0 | 898 | 1597 | 54546 | 29.27 | 0.41 |
| | 15 | 791 | 626 | 51610 | 12.13 | |

Table S3. Statistics used to calculate binding frequency in Supplementary Figure S2 and S4. $N_B$ refers to number of PNA binding events, $T_{tot}$ to total observation time, $f$ to binding frequency.

| **Construct** | **[PEG] (v/v) %** | **$N_B$** | **$T_{tot}$ (s)** | **f ($\times 10^{-3}$ $s^{-1}$)** |
|---|---|---|---|---|
| 1G-Tract (PEG-200) **Fig. S2** | 0 | 195 | 15025 | 12.98 |
| | 5 | 145 | 10677 | 13.58 |
| | 10 | 160 | 12232 | 13.08 |
| | 15 | 162 | 11637 | 13.92 |
| | 20 | 170 | 12890 | 13.19 |
| | 30 | 146 | 10950 | 13.33 |
| 1G-Tract (PEG-6000) **Fig. S2** | 0 | 292 | 25002 | 11.68 |
| | 5 | 310 | 28833 | 10.75 |
| | 10 | 265 | 23753 | 11.16 |
| | 15 | 182 | 16567 | 10.99 |
| | 20 | 170 | 14330 | 11.86 |
| | 30 | 150 | 13097 | 11.45 |
| 10T-1G-Tract (PEG-200) **Fig. S4** | 0 | 160 | 18951 | 8.44 |
| | 5 | 190 | 21475 | 8.85 |
| | 10 | 176 | 20015 | 8.79 |
| | 15 | 170 | 19975 | 8.51 |
| | 20 | 172 | 18985 | 9.06 |
| | 30 | 154 | 17805 | 8.65 |

### References


(1) Shiekh, S.; Mustafa, G.; Kodikara, S. G.; Hoque, M. E.; Yokie, E.; Portman, J. J.; Balci, H. Emerging Accessibility Patterns in Long Telomeric Overhangs. *Proc Natl Acad Sci U S A* 2022, *119* (30). https://doi.org/10.1073/pnas.2202317119.
